# Frontostriatal dynamics of cognitive control

**DOI:** 10.1101/2025.03.07.642117

**Authors:** Thomas W. Elston, Joni D. Wallis

## Abstract

A central tenet of cognitive psychology is the existence of two distinct modes of processing: the fast, automatic system and the slow, deliberative system ^1,2^. The prefrontal cortex is thought to control the balance between these two systems, allowing automatic processing where possible, but stepping in when deliberative processes are required, such as in novel, uncertain, or conflicting situations ^3–6^. Despite the central importance of this idea in cognitive neuroscience, there is little understanding of how this control process is implemented. We used Neuropixel recordings in rhesus macaques to decode neural activity and reveal the cognitive processes in prefrontal cortex and striatum during automatic and deliberative decisions. During automatic decisions, the chosen option was decoded simultaneously from both areas. During deliberative choices, the prefrontal cortex serially considered multiple options while neither could be decoded from the striatum. This suppression occurred via an increase in alpha band coherence between the two structures and scaled with the volatility of prefrontal representations. Our findings reveal a specific neural mechanism that may explain how frontostriatal circuits implement cognitive control and provide a novel mechanism by which to treat disorders of behavioral control.

## Introduction

A diverse array of cognitive processes, including attention ^1^, learning ^7,8^, and decision-making ^2,9^, have highlighted the existence of dual decision systems. The first system is algorithmically simple, operates quickly, but is inflexible, while the second system is computationally expensive, operates more slowly, but is better able to cope with changing demands, novelty, or complexity ^7,8^. These dual systems are thought to be instantiated in distinct neural circuits, with the striatum playing a central role in habitual, stimulus-driven responses, and the prefrontal cortex supporting flexible, goal-directed behavior ^4,7,10^. Electrophysiological and neuroimaging data support the involvement of prefrontal inhibition of the basal ganglia during cognitive control ^11–16^. However, these studies have been unable to resolve how the prefrontal cortex detects the need for control and how control is implemented because they could not capture the moment-by-moment dynamics through which prefrontal regions detect conflict and modulate the striatum. Recent advances in high-channel count recordings, particularly Neuropixel technology, coupled with decoding algorithms now enable measurement of cognitive processes with single-trial resolution ^22–24^. Here we use this approach to reveal how prefrontal volatility signals the need for control and how alpha-band synchronization implements this control by suppressing striatal activity.

## Results

We trained two monkeys (K and D) on a state-dependent decision task designed to evoke conflict between competing value schemas. We previously used a variant of this task to demonstrate the existence of state-dependent value codes in the orbitofrontal cortex (OFC) ^25^. On each trial, the monkeys were cued to use one of three value schemas corresponding to task states A, B, or C, each of which predicted different contingencies between a set of reward-predictive cues and reward amounts (**Fig. 1a**). After a brief delay, they were presented with either one (forced choice, 25% of trials) or two options (free choice, 75% of trials). In states A and B, the same options had different values depending on the state, while state C involved no such conflict. The subjects reported their choices by fixating on their chosen option for 300 ms.

**Figure 1.**
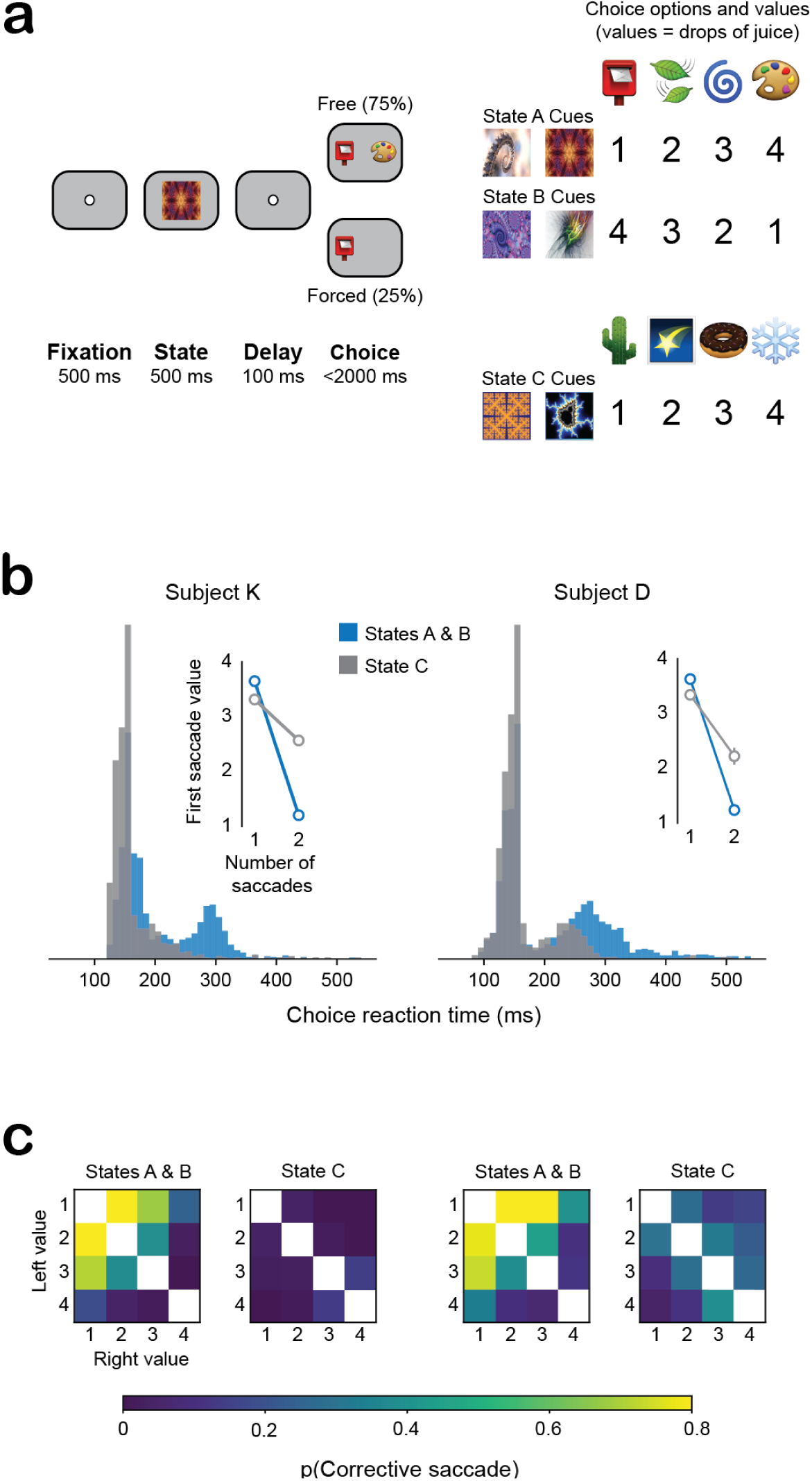
Behavioral task. (a) Subjects initially fixated and were then shown one of six state cues (two cues per state). After a delay, subjects were presented with either one or two choice options. The optimal choice depended on the state cued earlier in the trial. (b) Histograms of choice reaction times during each of the three states. The bimodal distribution in states A and B was due to the presence of corrective saccades. Inset: Mean (± 95% confidence intervals) value of the option selected by the subjects’ first saccade. When subjects made two saccades, the first saccade in states A and B (but not state C) was almost always to the the value 1 option (2-way ANOVA interaction with factors of number of saccades and task state, subject K: *F_1,2501_* = 150, *p* = 1.0 x 10^-33^, subject D: *F*_1, 2145_ = 140, *p* = 2.4 x 10^-30^). (c) Likelihood of corrective saccades in all choice conditions. Corrective saccades in states A and B were consistently associated with trials that contained the value 1 option, which would be the value 4 option in the alternative state.

The task design evoked behaviors in states A and B consistent with increased cognitive control. Most choices involved a single saccade to the chosen option, but on some trials the subjects first made a saccade to the unchosen option, resulting in a bimodal distribution of choice reaction times (**Fig. 1b**). Double saccades mainly occurred when value 1 was one of the options (**Fig. 1b, c**). Given that the best option (value 4) in state A was the worst option (value 1) in state B, these patterns reveal an initial automatic response toward options based on values from the alternative state, followed by top-down cognitive control that overrode this prepotent response. This control manifested as a second corrective saccade redirecting gaze to the optimal choice for the current state. The near-absence of such corrective saccades in state C, where no value conflict existed between states, further supports the interpretation that this behavior reflects an automatic pre-potent response being overridden.

To examine the neural mechanisms of top-down control, we simultaneously recorded from OFC and the caudate nucleus (CdN) of the striatum using Neuropixel probes (**Fig. 2a**). Across four recording sessions per subject, we obtained large numbers of well-isolated neurons from both regions (Subject K: 1528 OFC and 1085 CdN neurons; Subject D: 1293 OFC and 1149 CdN neurons; **Fig. 2b**). To quantify neuronal tuning, we performed a sliding 2-way ANOVA with factors of State (A, B, and C) and Value (1 thru 4). Neurons in OFC primarily encoded the state-value interaction (**Fig. 2c**), indicating neurons in this region encoded value in a state-dependent manner (**Fig. 2d**). In contrast, similar proportions of CdN neurons encoded value in either a state-dependent or state-independent way. State-dependent value encoding occurred significantly earlier in OFC than CdN (**Fig. 2e**).

**Figure 2.**
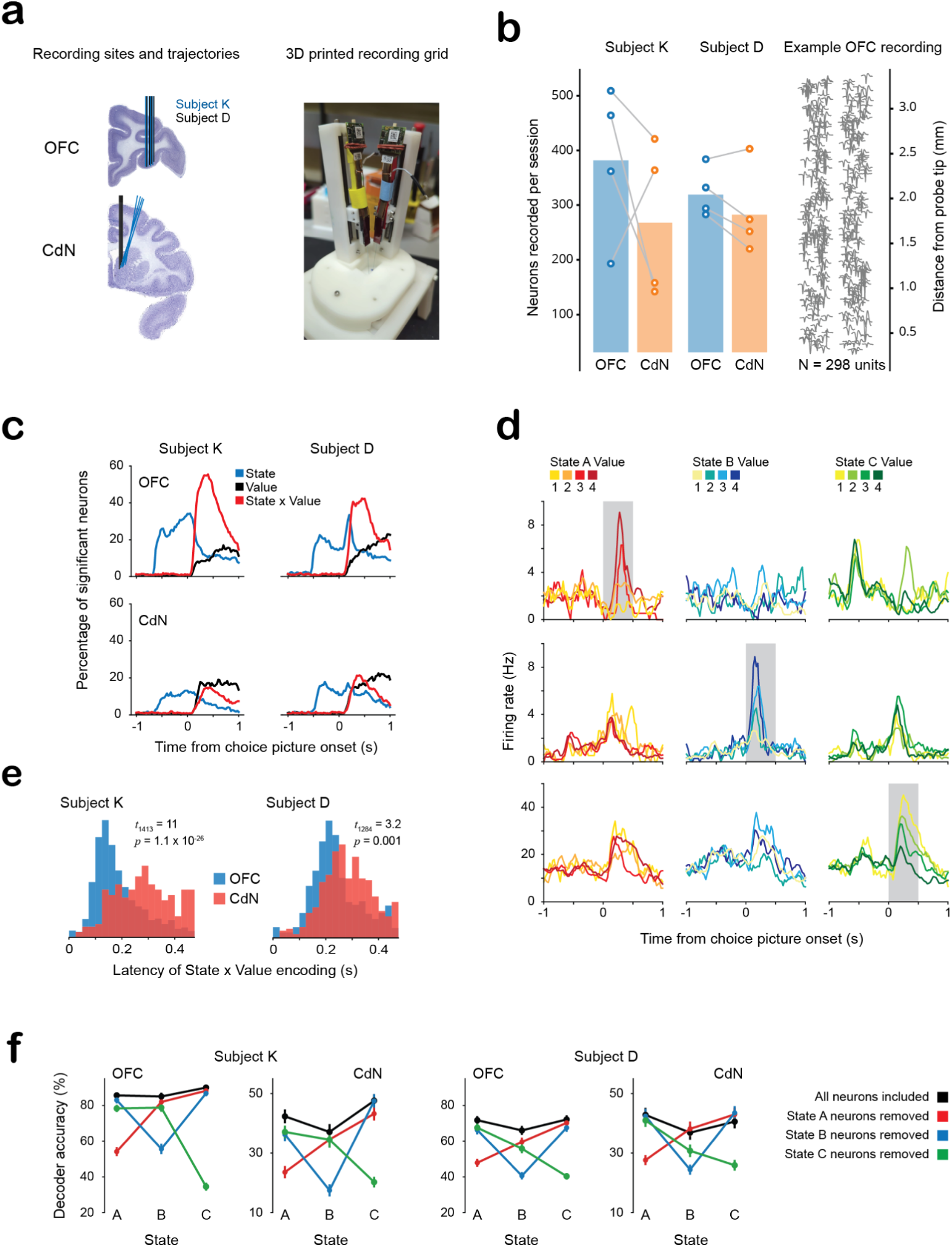
State-dependent value representations in OFC and CdN. (a) Electrode trajectories and 3D-printed recording grids for simultaneous neuropixel recordings in the OFC and CdN. There were four recordings per subject. (b) Numbers of simultaneously recorded neurons in each brain area and recording session. Linked OFC-CdN marker pairs indicate single sessions. Right: mean waveforms of 298 simultaneously recorded neurons in OFC localized onto the surface of the probe. Each trace denotes the mean waveform of a single isolated neuron. (c) Percentage of neurons significantly encoding State, Value, or their interaction via a sliding window 2-way ANOVA. During the choice epoch, OFC neurons encoded state-dependent values, whereas CdN neurons were evenly split between those encoding state-dependent and state-independent value information. (d) Three example OFC neurons that only encoded Value in State A (top row), B (middle row), or C (bottom row). The shaded region indicates the choice epoch. (e) State-dependent value encoding occurred significantly earlier in OFC before CdN in both subjects (unpaired *t*-tests). For subject D, OFC N = 859 state-value encoding neurons and CdN N = 427. For subject K, OFC N = 1151 CdN N = 264. (f) Population-level decoding also demonstrates the existence of distinct value ensembles in each state. drive decoder performance. Decoders were trained on all trials and the contribution of each ensemble of state-dependent value neurons was assessed by iteratively removing them and recalculating decoder performance. The change in decoder performance was largest for the state associated with the ensemble that was removed. Markers and error bars denote means and bootstrapped 95% confidence intervals. For subject D, state A N = 978 trials, state B = 829, and C = 1081 trials. For subject K, A = 1244, B = 935, and C = 1413.

Population-level decoding also revealed the presence of state-specific value ensembles in both OFC and CdN. We trained Linear Discriminant Analysis classifiers to decode specific combinations of state and value from simultaneously recorded neuronal activity (see Methods). For each brain area in each subject, we first established baseline decoder classification accuracy on single-saccade trials using the complete neuronal population. We then systematically set the decoder weights of each state-specific neuronal ensemble to zero and reassessed performance. The largest accuracy decrements occurred in states matching the removed ensemble (**Fig. 2f**). For example, when we removed the contribution of the neurons that encoded value in state A, decoder performance was selectively degraded during state A trials. This demonstrates that population-level state-dependent value representations rely on distinct ensembles in each brain area.

An advantage of population-level decoding is that it provides us with single-trial resolution of neuronal dynamics that can be used to determine whether there are differences in frontostriatal dynamics on trials where top-down control is or is not required. We focused on the trials from states A and B and compared dynamics when a corrective saccade was required versus trials where choice involved a single saccade. During single-saccade trials, both brain areas showed rapid, stable representation of the state-appropriate, high-value chosen option (**Fig. 3a**, left). However, during trials with a corrective saccade, OFC value representations vacillated between the state-appropriate value of the ultimately chosen option and the counterfactual value that the initially-chosen (but ultimately rejected) option would have had in the alternative state (**Fig. 3a**, right). For example, in the top-right panel of Fig 3a, we show a state A trial where Subject K chose between a postbox and blue-swirl. The monkey initially saccaded to the postbox (which had value 1 in the current state, state A, but would have had value 4 in state B) before correcting this selection and making a second saccade to choose the blue-swirl (value 3 in state A). During this trial, OFC activity fluctuated between encoding the postbox’s counterfactual value in state B (value 4) and the contextually appropriate value of the ultimately chosen blue-swirl (value 3). Such dynamics in CdN were much weaker and less frequent, suggesting that conflict-related volatility in the OFC may suppress value representations in the CdN. We confirmed that these dynamics were representative of the dynamics typically observed, by quantifying the frequency of these decoded representations across all state A and B trials (**Fig. 3b**).

**Figure 3.**
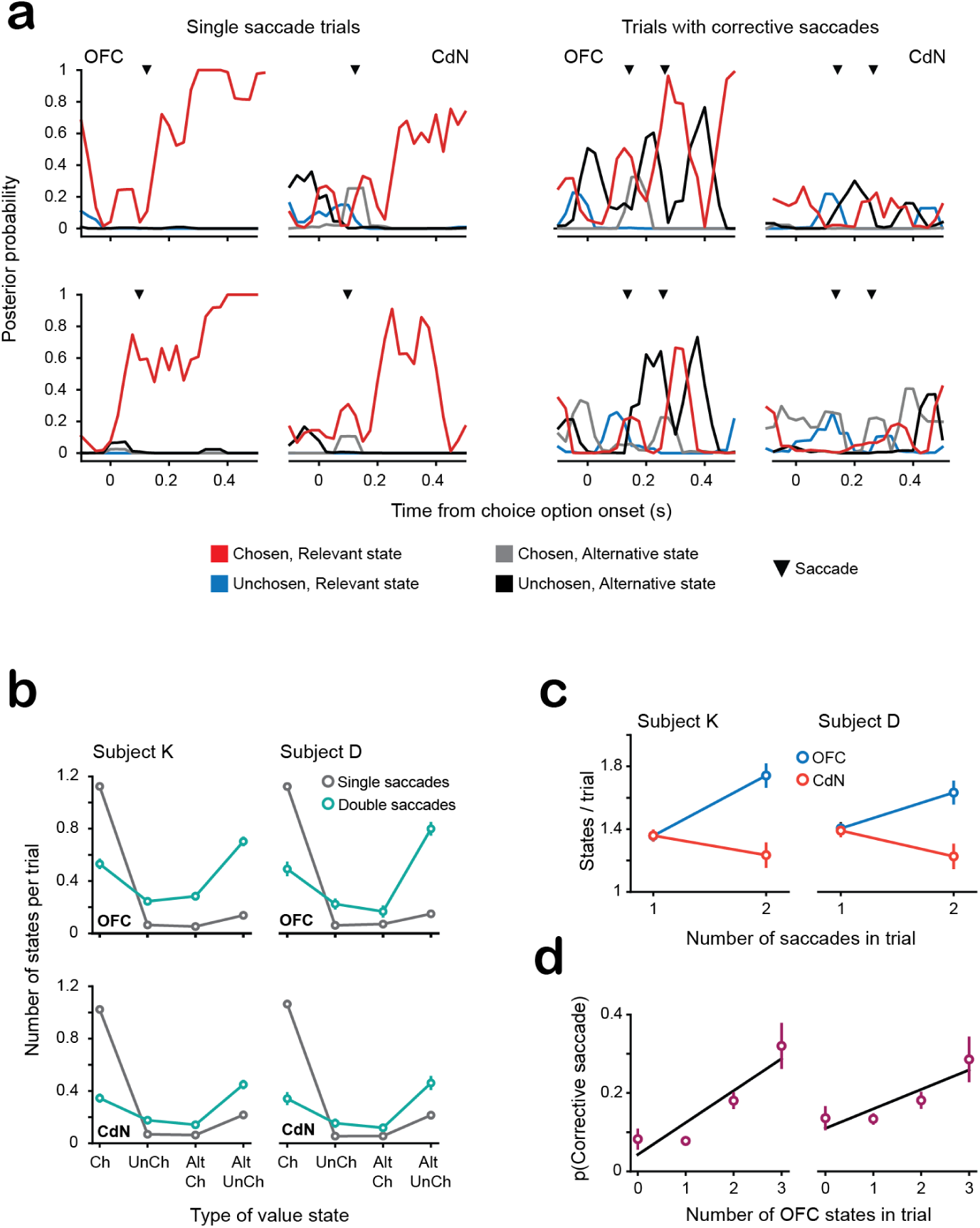
Fronto-striatal value dynamics during single and double saccade trials. (a) Decoding from example trials with either a single saccade (left) or two saccades (right). The triangular ticks indicate the onset of a saccade. During single saccade trials, both OFC and CdN value dynamics reflected the high value, chosen option. During trials with corrective saccades, OFC value dynamics vacillated between the state-appropriate, high value option and the counterfactual value associated with alternative state. These dynamics were not present in CdN. (b) Mean number of each task-relevant value state observed in a single trial. During single-take trials, both OFC and CdN value dynamics reflected the high value, ultimately chosen option. During trials with corrective saccades, the most commonly observed state was associated with the counterfactual value of the unchosen option (that is, the value of the unchosen option in the alternative task state). The second most commonly observed state was associated with the high value, contextually appropriate option that would ultimately be selected. Similar rankings were observed in CdN, although there were fewer states observed in general. For subject D, N = 1747 single saccade trials and 539 double saccade trials in states A and B. For subject K, N = 2137 single saccade trials and 546 double saccade trials in states A and B. The markers and error bars denote means and bootstrapped 95% confidence intervals. (c) Corrective saccades were associated with increased volatility in OFC and suppression of CdN. The y-axis shows the number of discrete value states observed in a single trial. During single saccade trials, a similar number of value states were observed in OFC and CdN. However, during double saccade trials, the number of states in OFC increased whereas the number of states decreased in CdN (2-way ANOVA interaction between the factors of brain area and number of saccades, subject K: *F*_2, 5370_ = 38, *p* = 3.9 x 10^-17^, subject D *F*_2, 4568_ = 26, *p* = 4.4 x 10^-7^). (d) Corrective saccades were significantly more likely when more value states occurred in OFC (logistic regression, subject K: log odds = 0.17, *p* = 1.2 x 10 ^-5^, subject D: log odds = 0.08, *p* = 0.003). Each datapoint denotes the mean and bootstrapped 95% confidence intervals.

To quantify the dynamics of the value coding, we measured the number of discrete value representations decoded during the choice epoch, the 500 ms period beginning from the onset of the choice options (**Fig. 3c**). During single saccade trials, the number of value states did not differ between OFC and CdN. However, when corrective saccades occurred, OFC showed a significant increase in the number of value states, while the number decreased in CdN. Furthermore, the number of distinct OFC value states significantly predicted the likelihood of a corrective saccade occurring (**Fig. 3d**).

These results suggest that top-down control is associated with increased volatility of OFC value representations induced by consideration of alternative contexts that results in a suppression of value representations in CdN. To determine the potential mechanism by which this is accomplished, we examined OFC-CdN local field potential synchronization (see Methods). During corrective saccades there was a significant increase in the synchronization of the alpha band between OFC and CdN relative to trials without corrective saccades (**Fig. 4a,b**). The effect was present in both subjects, although subject K had an additional peak in the beta band. Next, we assessed the directionality of OFC-CdN communication using a trial-by-trial amplitude cross-correlation analysis. Across both subjects and in each individual recording, we consistently found that OFC alpha activity preceded CdN alpha activity by approximately 10 ms for both single saccade trials and those with corrective saccades (**Fig. 4c**). In addition, there was a strong, positive relationship between the number of OFC value representations that were detected on a single trial and the strength of OFC-CdN alpha-band coherence (**Fig 4d**).

**Figure 4.**
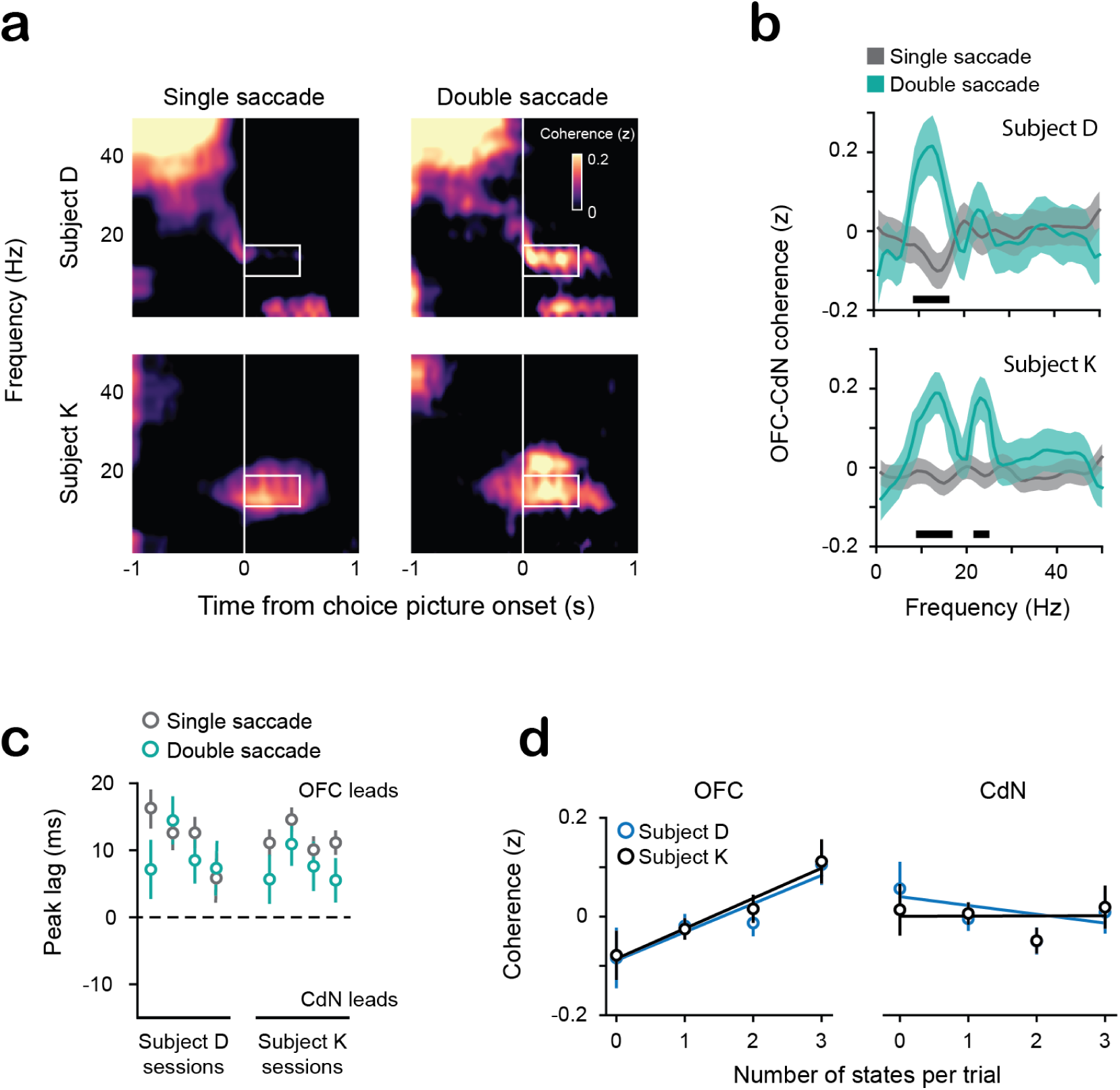
Fronto-striatal coherence during top-down control. (a) Spectral OFC-CdN coherence during single and double saccade trials of states A and B from a single session for subjects D (top) and K (bottom). The boxed region highlights that alpha-band coherence, centered at about 10 Hz, is strongly elevated during double saccade trials. (b) Quantification of OFC-CdN coherence during the choice epoch. Horizontal black bars denote significant differences (*p* < 0.001) at a specific frequency via unpaired *t*-tests. In subject D (top), the only significant difference was in the alpha band, at about 10 Hz. In subject K (bottom), significant differences were detected in both the alpha and beta bands. Number of trials as in Fig. 3b. The markers and error bars denote means and bootstrapped 95% confidence intervals. (c) OFC alpha consistently preceded CdN alpha. In all cases, the mean lag was significantly greater than zero (one-sample *t*-tests, *p* < 0.001), indicating OFC alpha activity preceded CdN alpha. For some sessions, there was a significant difference in the size of the lag on trials with single versus double saccades, but this was not consistent across sessions and was not significant when assessed across all trials in every session (two-sample *t*-test, subject D: *t*_1412_ = 1.3, *p* = 0.19, subject K: *t*_2003_ = 1.6, *p* = 0.12). The markers and error bars denote means and bootstrapped 95% confidence intervals for each session. (d) OFC volatility significantly predicted OFC-CdN alpha coherence, while there was no relationship with CdN volatility (2-way ANOVA with post hoc linear contrasts, significant interaction between factors of brain area and number of decoded states, subject D: *F*_1,7924_ = 7.3, *p* = 0.006; subject K: *F*_1, 8788_ = 11, *p* = 0.001). The relationship between the number of decoded states and alpha coherence were positive and significant in OFC (subject D: *β* = 0.057, *p* = 0.02, subject K: *β* = 0.061, *p* = 0.001) but not in CdN (subject D: *β* = −0.01, *p* = 0.33, subject K: *β* = 0.0008, *p* = 0.09). The markers and error bars denote means and bootstrapped 95% confidence intervals.

## Discussion

Prefrontal cortex has long been associated with inhibitory control, whereby more automatic responses are inhibited when they are not contextually relevant ^4,6,26^. Our findings reveal that during cognitive conflict, the OFC exhibits “volatility” – a rapid switching between multiple competing value representations rather than stably encoding a single value. This neural volatility in the OFC predicted both the likelihood of behavioral control (corrective saccades) and increased OFC-led communication with CdN in the alpha-band. Importantly, the strength of this alpha synchronization scaled directly with the degree of representational instability in OFC (measured as the number of distinct value states decoded within a trial) and was associated with suppressed value coding in CdN. This suggests that alpha-band synchronization may be a critical mechanism which enables the prefrontal cortex to exert top-down control over the striatum.

Alpha synchronization has previously been argued to be a mechanism of inhibition in sensorimotor regions ^27–29^. For example, neuronal firing rates in sensorimotor regions are suppressed at the peak of the alpha oscillation ^30^ and increased alpha activity is observed in occipital areas when ignoring visual distractors ^31^. Causal manipulations using MEG neurofeedback in humans biased visual attention away from the corresponding parietal areas in which alpha synchrony is high ^32^. The parallel between alpha-mediated inhibition in sensorimotor systems and our observed frontostriatal dynamics suggests that alpha synchronization may represent a general neural mechanism for implementing top-down control across neural systems.

Consistent with our previous results ^25^, OFC encoded value in a state-dependent manner. In contrast, CdN neurons exhibited both state-dependent and state-independent value coding. State-dependent value signals emerged significantly earlier in OFC than in CdN, consistent with previous evidence that striatal value signals are inherited from OFC ^33^. This temporal relationship, combined with the asymmetric distribution of state-dependent value coding across regions, suggests a hierarchical scheme where OFC maintains flexible, context-sensitive value representations that are used to inform value signals in CdN. Critically, this hierarchical arrangement provides a substrate for implementing cognitive control through the alpha-mediated top-down modulation of striatal activity that we observed during deliberative decision-making.

The striatum has long been implicated in habitual, stimulus-driven behavior ^10,18,34^, while prefrontal regions are critical for flexible, goal-directed actions ^4,7^. Disrupted connectivity between the prefrontal cortex and striatum has been identified as a key pathophysiological feature in several neuropsychiatric conditions characterized by cognitive inflexibility, including obsessive-compulsive disorder ^35,36^ and addiction ^37–39^. In conditions of cognitive inflexibility, patients often struggle to override habitual or prepotent striatal-mediated responses even when such responses become maladaptive. Our finding that precisely timed alpha-band synchronization between OFC and striatum enables successful corrective control suggests a specific neural mechanism that may be impaired in these disorders. The identification of alpha-band coherence as a specific marker of successful cognitive control could provide a useful physiological target for therapeutic interventions, such as closed-loop neurostimulation approaches ^40–42^.

## Methods

### Experimental model and subject details

All procedures were carried out as specified in the National Research Council guidelines and approved by the Animal Care and Use Committee at the University of California, Berkeley. Two male rhesus macaques (subjects D and K, respectively) aged 9 and 5 years, and weighing 12 and 12 kg at the time of recording were used in the current study. Subjects sat head-fixed in a primate chair (Crist Instrument, Hagerstown, MD). Eye movements were tracked with an infrared system (SR Research, Ottawa, Ontario, CN). Stimulus presentation and behavioral conditions were controlled using the MonkeyLogic toolbox ^43^. Subjects had unilateral recording chambers implanted that allowed access to both OFC and CdN.

### Behavioral task

Stimuli were presented on a computer monitor positioned at a viewing distance of approximately 30 cm. The subjects were trained to perform a state-dependent decision task. This required them to choose between eight reward-predictive pictures, where the reward amounts associated with the pictures depended on a cued task state (Fig. 1a). Subjects self-initiated trials by fixating on a white dot for 500 ms. After initial fixation, the subjects were shown a state cue that indicated whether the upcoming choice should be evaluated according to the value scheme associated with state A, B, or C. To unconfound neuronal activity related to the physical properties of the stimuli from the meaning they signified, we used two distinct cues for each state. Thus, we used two cues to indicate each state (six total state cues). After 500 ms, the state cue disappeared, and the subjects had to maintain central fixation for a further 100 ms. Finally, the choice options were presented. On 75% of the trials two options were shown (free choice trials), while on the remaining 25% of the trials there was only one option (forced choice trials). The inclusion of forced choice trials ensured that subjects regularly experienced all reward contingencies. Subjects reported their response by shifting their gaze to their selection and fixating for 300 ms. The values of the choice options corresponded to different amounts of apple juice reward.

To illustrate how state modulates value and choice, consider a trial where the state cue was a tentacle-like fractal (indicating state A), and the choice options were the blue swirl and postbox emojis. In state A, selecting the blue swirl yielded three drops of juice whereas selecting the postbox yielded only one drop of juice. However, had a state B cue been shown earlier (e.g., an orange fractal), selecting the postbox would have yielded four drops of juice and the blue swirl only two drops. State cues were pseudorandomized across trials. State C functioned as a control to assess neural responses during choice where cognitive control was not required. It had a different set of choice options to states A and B and there was only one value schema.

### Neurophysiological recordings

Subjects were fitted with titanium head positioners and imaged in a 3T magnetic resonance imaging scanner. The resulting images were used to generate 3D reconstructions of each subject’s skull and brain areas of interest. We then implanted custom, radiotranslucent recording chambers made of polyether-ether-ketone (PEEK; Gray Matter Research, Bozeman, MT). Each session we acutely lowered two 45 mm primate Neuropixels probes (IMEC VZW, Leuven, Belgium) into area 13 of OFC and the ventromedial aspect of CdN, which is a major striatal projection target of OFC ^44^. Electrode trajectories were determined by incorporating the subject’s MRI into computer-assisted drawing software (OnShape, Cambridge, MA) to design and 3D print custom recording grids to lower the probes (Form 4 3D printer, Form Labs, Somerville, MA). After insertion, the probes were allowed to settle for around 60 min before the start of the experiment, to mitigate drift. We configured probes to record from 384 active channels in a contiguous block at the tip, allowing dense sampling of neuronal activity along a 3.84 mm span. Neuronal activity was filtered and digitized for action potential bands (300 Hz high-pass filter, 30 kHz sampling frequency) and local field potential (1 kHz low-pass filter, 2.5 kHz sampling frequency). Activity was monitored during experimental sessions and saved to disk using SpikeGLX (https://billkarsh.github.io/SpikeGLX/).

Spiking in the action-potential band was identified and sorted offline using *Kilosort4* ^45^. Because each Neuropixels headstage samples the data with a slightly different system clock, it was necessary to map all physiological and task-event time series into a common timeline. We accomplished this by broadcasting a 5 Volt, 1 Hz square wave to a unique reference channel on the Neuropixel probes (channel 385) and the non-neural acquisition board. We detected and stored the times of the rising phase of the 5 V wave on each cycle and then used regression to interpolate all events into the timeline of the first Neuropixels probe. To be included for further analysis, neurons had to be present for >90% of the experimental session, have a mean firing rate over the course of the entire session >1 Hz, and <0.1% of spikes occurring within 1 ms of another isolated neuron’s spikes (that is, <0.1% interspike interval violations).

### Single-neuron tuning

To quantify the tuning of single neurons and its dynamics, we used a sliding-window two-way ANOVA with factors of Task (states A, B, and C), and Value. Factors were dummy coded and centered in order to ensure independence of the interaction term. Windows were 100 ms long and stepped by 25 ms. Significance was assessed at *p* < 0.01.

To better understand state-value interactions, we first identified neurons that encoded significant state-dependent values. For each neuron, we calculated its mean firing rate, *F*, during the choice epoch of single-saccade trials. We then performed a two-way ANOVA with factors of Task (states A, B, and C), and Value. For neurons that encoded a significant state-value interaction we conducted the following regression:

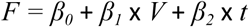

where *V* indicated the number of juice drops associated with the chosen option and *t* was a nuisance parameter that indicated the trial number and was used to account for non-stationarity in the neuronal recordings. We performed this analysis separately for trials in state A, B, or C and defined a neuron’s preferred state as the one with the largest *V* coefficient.

### Population-level decoding of value information

We trained Linear Discriminant Analysis (LDA) decoders to predict state-dependent values from OFC and CdN neural activity. For each brain area, the decoder classified 12 distinct value classes (4 possible reward values × 3 task states) using z-scored firing rates from hundreds of simultaneously recorded neurons. In each of 1000 bootstrap iterations, we randomly selected 90% of single-saccade trials for training and used all remaining trials for testing. The decoder computed posterior probabilities for each value class in the test trials. This procedure was repeated across time using 100 ms bins with a 25 ms step between bins. Across bootstraps, all trials were used in the testing set and we averaged across bootstrap iterations where the same trial appeared in the test set. Thus, we obtained trial-by-trial estimates of the relative strength of each state-value at each timestep for both brain areas. For each trial, we then recategorized the decoded values based on their behavioral relevance: whether they corresponded to the chosen or unchosen option, and whether they represented the value in the current task state or the alternative state (for states A and B, where stimulus values differed by task state).

To evaluate how state-specific value ensembles might contribute to population dynamics, we implemented a decoder lesioning procedure. Similar to our primary decoding analysis, we used a bootstrapping approach (1000 iterations) on single-saccade trials, but focused exclusively on neural activity during the 500 ms period following choice option onset. After establishing baseline classification accuracy across all task states using the complete neuronal population in each area, we systematically lesioned state-specific ensembles (neurons showing selective value encoding in states A, B, or C) by setting their corresponding decoder weights to zero. We performed this procedure separately for the state A, B, and C ensemble. While removing neurons from the decoder was expected to reduce overall accuracy, we predicted that the largest performance decline would occur in the task state corresponding to the lesioned ensemble.

To assess the stability of value representations during decision-making, we identified sustained periods of confident value decoding, defined as the posterior probability for a given value to exceed twice the chance level (16.7%, or 2/12) for at least two consecutive time bins (50 ms). These thresholds were selected because they yielded a false positive rate during the fixation epoch below 5% across all 12 value classes. We then quantified the number of discrete value representations decoded during the choice epoch.

### Local field potential analysis

We computed spectral coherence between simultaneously recorded OFC and CdN LFPs using the scipy.signal.coherence method in Python. Spectral coherence quantifies the frequency-dependent correlation between two signals, measuring how consistently the phase relationships between frequency components are maintained over time, normalized by the power in each signal. This normalization by power means that coherence measures the consistency of phase relationships independent of signal amplitude, with values ranging from 0 (no phase synchronization) to 1 (perfect phase synchronization) at each frequency. Our parameters were a sampling frequency of 1000 Hz, a 1 second reading window with 95% overlap between windows, and a Hamming window as the tapering function. We computed the OFC-CdN coherence at each timestep of each trial. For statistical comparisons between single and double-saccade trials, we focused on the 500 ms period following choice option onset, comparing coherence values at each frequency using unpaired *t*-tests.

Because our spectral coherence analyses revealed significant synchronization specifically in the alpha band, we focused our analysis of signal directionality on this frequency range (8-16 Hz). We calculated the amplitude cross-correlation. We selected this analysis method over other measures such as Granger Causality or partial directed coherence, because it has been found to be as sensitive as these to measures to true changes in directionality while being less prone to producing false-positive results ^46^. For each trial, we first band-pass filtered both signals in the alpha band (8-16 Hz) using a zero-phase Butterworth filter. We then extracted the absolute analytic amplitude of these filtered signals using the Hilbert transform and computed cross-correlations between these amplitude envelopes with lags ranging from −100 to 100 ms. For each trial, we identified the lag that produced the maximum cross-correlation value. Positive lags indicate OFC activity preceding CdN, while negative lags indicate CdN leading OFC. We compared the distribution of optimal lags between single saccade trials and those with corrective saccades using unpaired *t*-tests, and tested whether the mean lag in each session differed significantly from zero via one-sample *t*-tests.

